# Fluorescence anisotropy structured illumination microscopy for quantitative super-resolved mapping of cell microenvironment and cytoskeletal dynamics

**DOI:** 10.64898/2026.03.06.710005

**Authors:** Shu Gao, Wenyi Wang, Liang Qiao, Han Wang, Mingqiao Liu, Yiwei Hou, Guangwei Xin, Chunyan Shan, Donghyun Kim, Zhixing Chen, Meiqi Li, Peng Xi

**Author notes:** Contributed equally to this work (Shu Gao, Wenyi Wang, Liang Qiao).

## Abstract

The crowded intracellular milieu shapes organelle architecture and dynamics, yet nanoscale heterogeneity in its physicochemical properties remains difficult to visualize with conventional fluorescence anisotropy (FA) imaging. Here, we develop fluorescence anisotropy structured illumination microscopy (FA-SIM), which employs orthogonal-polarization structured illumination with dual-angle detection to achieve ∼100-nm resolution and quantitative FA retrieval with 0.56% relative error, representing over 20-fold higher accuracy than conventional FA imaging. With low phototoxicity, FA-SIM enables dual-color, hour-long quantitative super-resolution imaging in cells. Using viscosity standards, defined nanoparticles and small-molecule drug binding assays, we validate FA-SIM as a quantitative reporter of rotational mobility and molecular interactions. In cells, FA-SIM resolves nanoscale crowding heterogeneity, correlates anisotropy landscapes with condensate dynamics, and uncovers a radial crowding gradient across the microtubule network and mitotic spindle. Long-term dual-color imaging further resolves coordinated actin–microtubule remodeling and associated microenvironmental changes. By enabling quantitative, super-resolved mapping of intracellular physical properties in living systems, FA-SIM provides a powerful platform for investigating the physical regulation of cellular organization and dynamics in health and disease.

## Introduction

The interior of living cells constitutes a highly complex physicochemical environment, where macromolecules such as ribosomes, proteins, and nucleic acids share a significant fraction of the available space and create a highly crowded and viscous milieu known as macromolecular crowding^1–6^. These properties significantly influences a wide array of cellular processes, including phase separation^2, 7–9^, DNA replication^10, 11^, protein folding^12–16^, cytoskeletal dynamics^17–20^, and intracellular cargo transport^21, 22^. Consequently, quantification of the cellular microenvironmental properties at high spatial and temporal resolution is crucial for a mechanistic understanding of cellular homeostasis and function.

Fluorescence anisotropy (FA) imaging provides a quantitative and noninvasive approach to probe such properties^23–26^. Defined as the extent of fluorophore rotational diffusion during its excited-state lifetime, anisotropy is sensitive to viscosity, temperature, and molecular volume, thereby reporting the local physical state of the environment. However, the application of FA microscopy is hindered by the lack of commercial systems. The researchers have to use polarization spectrophotometers to perform such analysis, which completely lose the spatial information. Moreover, conventional FA microscopies are often home-built and typically implemented under widefield illumination, suffering from inherently limited spatial resolution, noises from the out-of-focus layer, and compromised quantitative accuracy. While confocal^27^, selective plane illumination microscopy (SPIM)^28^ and two-photon exciation^29^ approaches offer optical sectioning capability, the spatial resolution of these modalities is fundamentally limited by diffraction, thus inadequate for capturing nanoscale microenvironmental dynamics in living systems. Consequently, a method capable of delivering super-resolution, quantitatively accurate FA imaging under live-cell conditions has remained an unmet need.

Structured illumination microscopy (SIM) provides an elegant solution to these challenges. SIM combines low phototoxicity with twofold improvement in spatial resolution and is inherently compatible with polarization-resolved excitation, as it used linearly polarization illumination patterns^30–34^. However, integrating FA with SIM requires overcoming multiple technical barriers, including precise polarization alignment, suppression of out-of-focus anisotropy contamination and quantitative image reconstruction for FA measurement.

Here, we develop a fluorescence anisotropy structured illumination microscopy (FA-SIM) imaging system that integrates orthogonal structured illumination with polarization-resolved detection and quantitative reconstruction algorithms. This system leverages the optical sectioning (OS) capability of structured illumination to suppress out-of-focus fluorescence and employs orthogonal-polarization excitation for cross-validation of anisotropy measurements, thereby enhancing both spatial resolution and quantitative precision. Combined with our quantitative image reconstruction algorithm, FA-SIM system achieves 100-nm lateral resolution and 0.56% relative error in anisotropy retrieval, under phototoxicity levels low enough for long-term live-cell imaging. Using this platform, we quantitatively map viscosity, rotational mobility, molecular binding and macromolecular crowding at nanoscale resolution, and visualize dynamic changes in cytoskeletal organization and intracellular physicochemical environments in real time. Together, FA-SIM provides a functional super-resolution approach for quantitatively mapping the physicochemical landscape of the living cell.

## RESULTS

### Development and calibration of the FA-SIM imaging system

To develop the FA-SIM imaging system, we used two orthogonal polarization angles (H and V) for the SIM pattern generation, ensuring the *s*-polarization illumination to minimize the axial excitation. The polarization properties of the excitation light were evaluated, and the high degree of linear polarization ensures accurate fluorescence anisotropy measurements and high-quality image reconstruction (Supplementary Table 1). The detection channels contain two orthogonal polarizations (H and V) as well, thus they serve as parallel and perpendicular polarization channels respectively when interlacing the excitation polarizations (Fig. 1a, b, Extended Data Fig. 1a, b). The geometric angle between the excitation and detection were pre-aligned to ensure the accuracy. The OS capability of the structured illumination microscopy effectively eliminates out-of-focus background signals^35^ to ensure only the in-focus plane FA imaging. Furthermore, the orthogonal angle structured illumination enables cross-validation of the distribution of molecules, making it insensitive to the polarization nature of the specimen itself, thereby enhancing the accuracy of fluorescence anisotropy measurements. Leveraging the super-resolution capability of SIM, our FA-SIM platform enables super-resolution florescence anisotropy imaging and quantification.

**Fig. 1.**
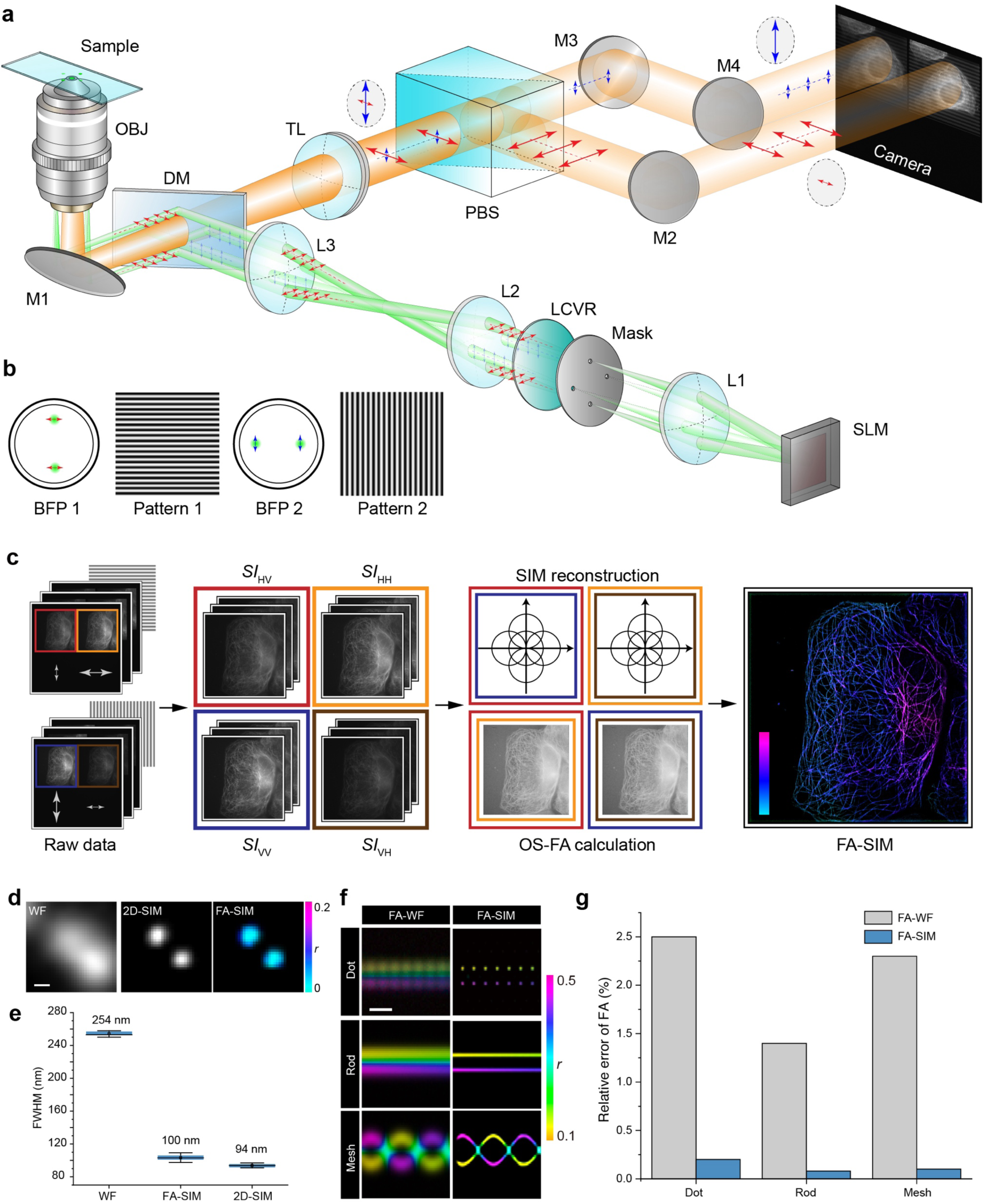
Design and performance characterization of the FA-SIM system. **a,** Schematic of the optical setup of FA-SIM system. SLM: spatial light modulator; LCVR: liquid crystal variable retarders; OBJ: objective lens; DM: dichroic mirror; TL: tube lens; PBS: polarization beam splitter; M1–4: mirror; L1–3: lenses. The arrow indicates the direction of light polarization. The red and blue arrows indicate orthogonal polarization directions. **b,** Schematic diagram of the diffraction pattern at the back focal plane (BFP) of the objective, polarization orientation, and structured illumination pattern for two-direction excitation. **c,** Image reconstruction workflow of FA-SIM imaging system. Raw images (*n* = 6) are divided into four groups (*SI*_HV_, *SI*_HH_, *SI*_VV_, *SI*_VH_), where the first symbol denotes the polarization of the illumination light and the second indicates the detected emission polarization. Each group contains three phase-shifted images. SIM reconstruction and fluorescence anisotropy calculations are performed to generate the final FA-SIM image. **d,** Representative imaging results of beads with the diameter of 40-nm using different imaging modalities, including WF, 2D-SIM and FA-SIM. Scale bar, 100 nm. **e,** Box chart of the measured FWHM values in different imaging modalities. *n* = 100 beads for statistic calculation. Error bars indicate mean ± S.D. **f,** Simulated FA-WF and FA-SIM images of dot-like, rod-like and mesh-like structures. Scale bar, 500 nm. **g,** Relative error of FA (%) in FA-WF and FA-SIM images in **(f)**.

For FA-SIM image reconstruction, a total of six raw images were acquired, comprising three phases for each of two orthogonal illumination directions. These images were grouped into four categories based on illumination and detection polarization: *SI*_HH_, *SI*_HV_, *SI*_VV_ and *SI*_VH_. After image registration and optical sectioning to eliminate out-of-focus signals, fluorescence anisotropy was calculated from the *SI*_HH_/*SI*_HV_ and *SI*_VV_/*SI*_VH_ image pairs using the formula defined as:

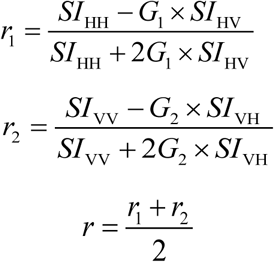

where *G*_1_ and *G*_2_ denote the polarization correction factors (Extended Data Fig. 1c, d).

The resolved anisotropy value *r* was used as the color-dimension for the proceeding SIM reconstruction (*r* matrix and the SIM super-resolution images are used in the Hue and Value channel of the FA-SIM final results, respectively, forming a HSV color space) (Fig. 1c, Supplementary Note 1).

Next, we characterized the performance of our FA-SIM system. Fourier analysis showed that FA-SIM effectively extended the accessible spatial frequency range relative to WF microscopy (Extended Data Fig. 2a, b). Imaging of 40-nm fluorescent beads revealed a marked improvement in lateral resolution compared with WF microscopy, enabling FA-SIM to resolve adjacent beads that were indistinguishable in WF images while simultaneously providing anisotropy information beyond conventional 2D-SIM. Gaussian fitting showed full widths at half maximum (FWHM) of 100 nm for FA-SIM and 254 nm for WF imaging (Fig. 1d, e). Consistently, decorrelation analysis yielded lateral resolutions of 96.4 nm for FA-SIM and 230 nm for WF (Extended Data Fig. 2c–e). Thus, FA-SIM achieves ∼2.5-fold improvement in resolution over WF microscopy.

To assess the quantitative accuracy of FA measurements in FA-SIM, we first examined the linearity of fluorescence intensities in the optically sectioned SIM pattern. The sectioned images exhibited a linear correlation coefficient of 0.88 with WF intensities, indicating that OS effectively suppresses out-of-focus background without introducing nonlinear intensity distortions, thereby preserving the validity of quantitative FA computation (Extended Data Fig. 2f, g). We next simulated a 3D Siemens star phantom to compare FA quantification in WF and structured-illumination modes. FA-SIM effectively removed the out-of-focus background signal, with enhanced spatially resolution and imaging quality (Extended Data Fig. 2h). Relative errors in FA retrieval were reduced from 12.3% in FA-WF to 0.89% in FA-SIM, and further to 0.56% when incorporating orthogonal-pattern cross-validation, constituting > 20-fold accuracy improvement in total (Extended Data Fig. 2i, j). FA-SIM also performed robustly across dot-like, rod-like and mesh-like test structures, resolving finer features and yielding more accurate FA quantification than WF imaging (Fig. 1f, g).

Together, these results demonstrate that FA-SIM substantially enhances both spatial resolution and quantitative fidelity in FA imaging, providing a powerful tool for nanoscale quantitative visualization of the intracellular microenvironment in cells.

### FA-SIM quantifies viscosity and target engagement in *vitro* and in *vivo*

Having established and calibrated the FA-SIM imaging system, we next sought to validate its functional proficiency in measuring molecular-scale properties through fluorescence anisotropy. We first assessed its sensitivity to microenvironmental viscosity using fluorescein in glycerol solutions with different viscosity. As expected, the FA-SIM images revealed a homogeneous and viscosity-dependent increase in anisotropy, with quantified values fitting the predicted Perrin relationship, confirming that our platform can accurately map viscosity variations at a large range (Fig. 2a, b).

**Fig. 2.**
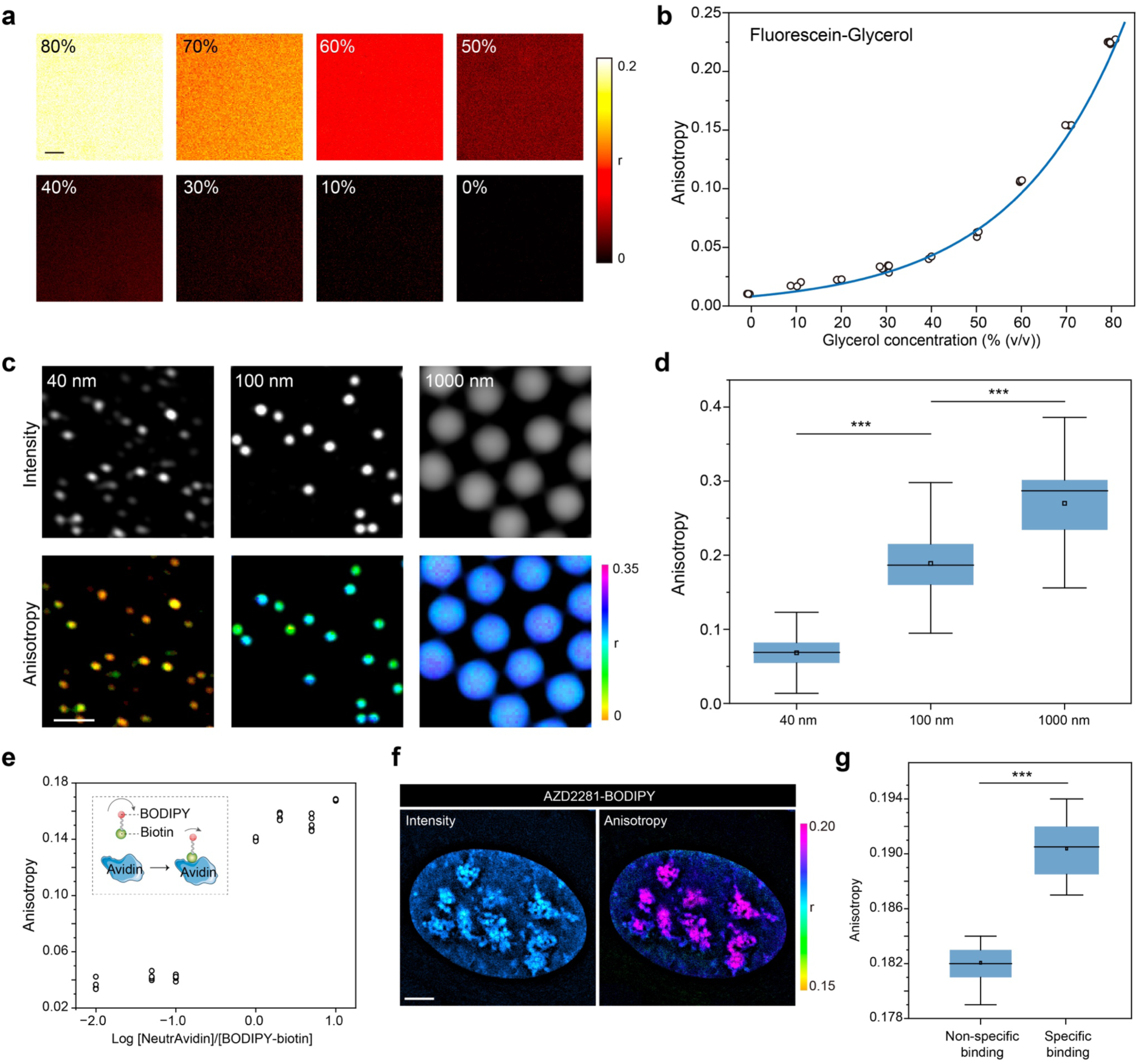
FA-SIM imaging indicates viscosity changes and target engagement. **a,** Fluorescence anisotropy images of fluorescein in glycerol solutions with increasing glycerol concentrations (v/v %), demonstrating sensitivity to changes in local viscosity. **b,** Quantification of fluorescence anisotropy from the images in **(a)**. Data points (open circles) represent individual measurements (*N* = 5), the blue line indicates the fitted curve. **c,** Representative fluorescence intensity (upper) and anisotropy (lower) images of fluorescent beads with diameters of 40-nm, 100-nm and 1000-nm. **d,** Quantitative comparison of fluorescence anisotropy values for the 40-nm, 100-nm and 1000-nm beads shown in **(c)**. *n* = 100 beads in each group for statistic comparison. **e,** Increase in fluorescence anisotropy of 1 μM BODIPY-biotin (MW 0.67 kDa) upon titration with NeutrAvidin (MW 60 kDa), indicating molecular binding and complex formation. Individual data points (open circles) are shown (*N* = 5). **f,** Representative fluorescence intensity (left) and anisotropy (right) images of U2OS cells treated with 1 μM AZD2281-BODIPY for 1 min at 37℃. **g,** Quantitative comparison of FA values of non-specific and specific bound drug. (*n* = 20 cells). Error bars and error bands indicate mean ± S.D. One-way ANOVA **(d)** and Two-tailed T-test **(g)** for the statistic calculation. *p* < 0.001: ***. Scale bars, 10 μm in **(a)**, 1 μm in **(c)** and 2 μm **(f)**.

To further demonstrate the capability of FA-SIM imaging system for resolving the rotational volumes, we imaged Alexa Fluor 488-labeled fluorescent beads with diameters of 40-nm, 100-nm and 1000-nm. FA-SIM images confirmed the super-resolution performance of our system and, crucially, revealed clear size-dependent rotational diffusion. Beads of different diameters exhibited distinct anisotropy signatures, with smaller beads displaying markedly lower FA values, consistent with their faster rotational dynamics (Fig. 2c, d).

We next applied our FA-SIM platform to monitor target engagement within biological systems. In a purified assay, the anisotropy of the small-molecule probe BODIPY-biotin (0.67 kDa) increased progressively upon binding to its high-molecular-weight target NeutriAvidin (60 kDa), verifying the detection of biomolecular interactions (Fig. 2e, Supplementary Note 2). In living U2OS cells, the BODIPY-conjugated PARP inhibitor AZD2281 showed rapid and specific nuclear accumulation within one minute of treatment (Fig. 2f). FA-SIM imaging provided a direct visual readout of target engagement, revealing distinct subnuclear structures with elevated anisotropy that reflected specific binding of AZD2281 (Fig. 2g).

Together, these data prove FA-SIM as a quantitative, super-resolution approach for mapping physicochemical microenvironments and molecular interactions in living cells.

### Mapping cytoplasmic macromolecular crowding at nanoscale resolution

Next, we extended FA-SIM to quantify the macromolecular crowding within living cells, a physical property that plays a central role in the densely packed intracellular space^1, 5^. To establish a quantitative readout for crowding, we used enhanced green fluorescence protein (EGFP) as a genetically encoded anisotropy sensor^36^. *In vitro*, the fluorescence anisotropy of EGFP increased monotonically with rising concentrations of bovine serum albumin (BSA) and Ficoll 400, used as crowding agents, demonstrating sensitivity to excluded-volume effects (Fig. 3a, Extended Data Fig. 3a). By contrast, EGFP anisotropy remained constant in NaCl solutions of varying concentration (Extended Data Fig. 3b), confirming that the response reflects macromolecular crowding rather than changes in ionic strength.

**Fig. 3.**
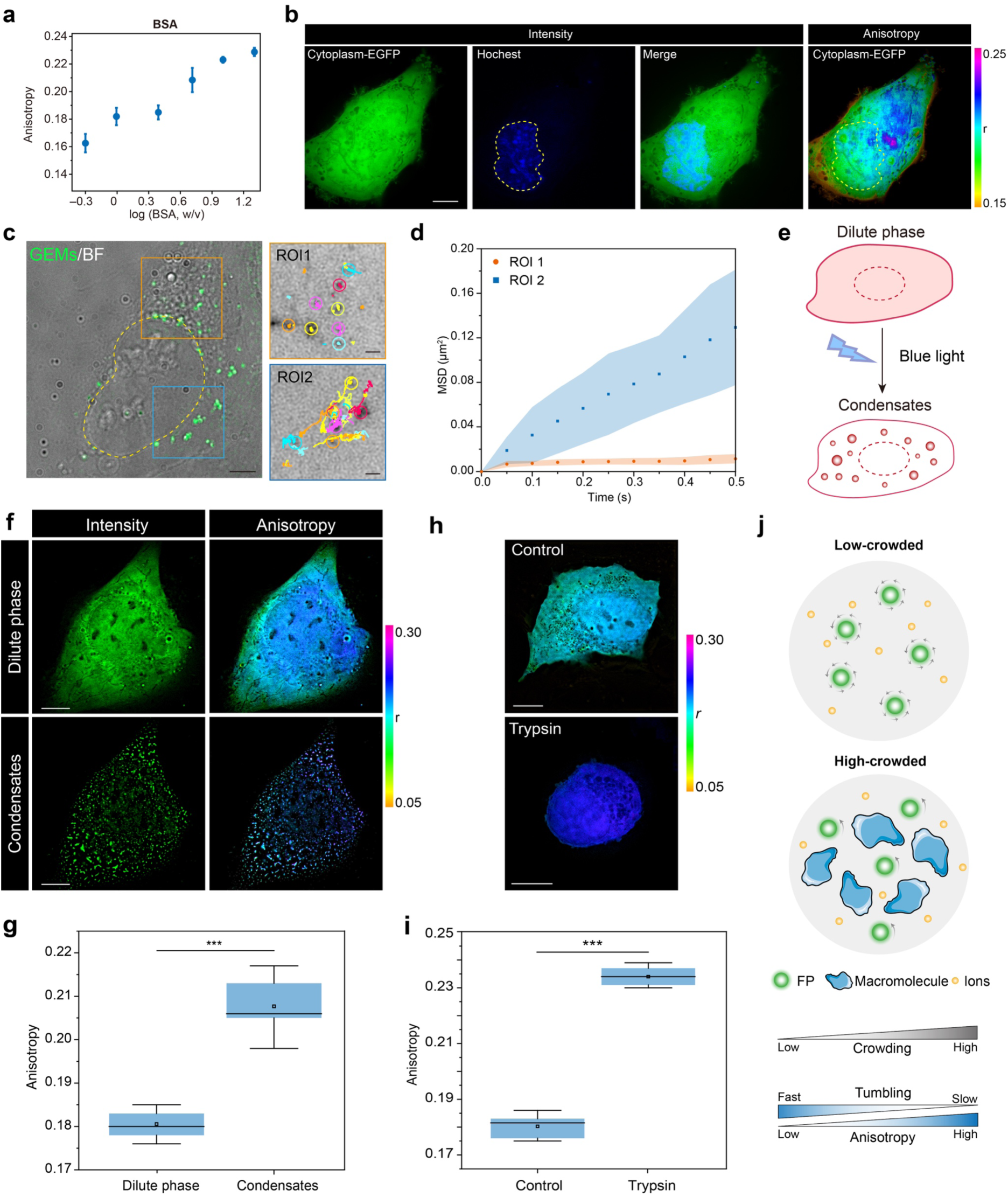
Probing macromolecular crowding via fluorescence protein anisotropy. **a,** Fluorescence anisotropy of EGFP in BSA solutions at varying concentrations. *N* = 5 biological replicates. **b,** FA-SIM imaging of a U2OS cell with cytoplasmic EGFP expression. Left: fluorescence intensity image (EGFP, green; nuclei, blue). Right: fluorescence anisotropy map; yellow dashed lines indicating nuclear contours. **c,** Left: U2OS cells expressing 40 nm genetically encoded multimeric nanoparticles (GEMs, green) and corresponding bright-filed image (gray). Scale bar: 10 μm. Right: time-lapse trajectories of GEMs (20 fps acquisition over 2 minutes); each color represents a single particle trajectory. Scale bar: 500 nm. **d,** Mean squared displacement (MSD) analysis of GEMs in organelle-dense (ROI 1) and organelle-sparse (ROI 2) regions (*n* = 20 cells). **e,** Schematic of light-induced phase separation in U2OS cells transfected with ESR1 IDR-mCherry-CRY2 plasmid. **f,** Fluorescence intensity (left) and fluorescence anisotropy (right) images of a U2OS cell before (upper) and after (bottom) light-induced phase separation. **g,** Comparison of mean fluorescence anisotropy of mCherry in U2OS cells before (Dilute phase) and after (Condensates) light-induced phase separation (*n* = 20 cells per group). **h,** Representative FA-SIM images of cytoplasmic EGFP in control and trypsin-treated U2OS cells. **i,** Comparison of mean fluorescence anisotropy of EGFP in control and trypsin-treated U2OS cells (*n* = 20 cells per group). **j,** Schematic model of EGFP as a sensor of macromolecular crowding. Error bars and error bands indicate mean ± S.D. Two-tailed T-test for the statistic calculation. *p* < 0.001: ***. Scale bars: 10 μm in (**b, f** and **h**).

To assess intracellular crowding in living cells, we expressed EGFP in the cytoplasm of U2OS cells for FA-SIM imaging. While fluorescence intensity appeared uniformly distributed, anisotropy maps revealed significant nanoscale heterogeneity in molecular mobility. Elevated anisotropy was consistently detected in the organelle-rich perinuclear region compared with the cell periphery (Fig. 3b), a pattern replicated with the fluorescent proteins mScarlet and mCherry (Extended Data Fig. 3c, d). These spatial variations are consistent with local differences in crowding density, where higher anisotropy reflects more restricted rotational diffusion.

To independently validate that anisotropy variations report genuine differences in cytoplasmic crowding, we tracked the motion of 40-nm genetically encoded multimeric nanoparticles (GEMs), an established reporter of intracellular density^2^. Bright-field imaging identified two characteristic regions: an organelle-dense perinuclear area (ROI 1) and an organelle-sparse peripheral area (ROI 2). Particle tracking revealed significantly constrained GEMs mobility in ROI 1, consistent with higher local crowding (Fig. 3c, Supplementary Note 3). Quantitative mean square displacement (MSD) analysis further confirmed substantially reduced mobility in perinuclear regions, independently supporting the anisotropy-derived crowding gradient (Fig. 3d). We further corroborated these findings using FRAP experiment. EGFP was ubiquitously expressed in U2OS cells, and regions at varying distances from the nucleus were photobleached under uniform laser power. Fluorescence recovery curves showed that EGFP near the nucleus exhibited the slowest recovery rate and lowest mobile fraction, whereas the most peripheral regions displayed the fastest and most complete recovery (Extended Data Fig. 3e, f). Together, these measurements confirm elevated macromolecular crowding and viscosity in the perinuclear cytoplasm.

Because macromolecular crowding plays a key role in regulating biomolecular condensation^20, 37–39^, we next examined liquid-liquid phase separation using FA-SIM. U2OS cells expressing ESR1 IDR-mCherry-CRY2 were stimulated with blue light to induce phase separation and condensate formation (Fig. 3e, Supplementary Video 1). Before induction, mCherry displayed a diffuse distribution with an anisotropy pattern similar to EGFP, yielding a mean anisotropy of 0.180. Upon stimulation, mCherry concentrated into condensed liquid droplets, and the increased crowding within droplets restricted molecular mobility, raising the mean anisotropy to 0.207 (Fig. 3f, g). Tracking the motion of condensates after phase separation revealed spatial dynamics that directly mirrored the pre-existing anisotropy landscape. Liquid droplets in the perinuclear region, where anisotropy was higher before induction, exhibited markedly slower movement, consistent with elevated crowding. By contrast, droplets in peripheral regions, characterized by lower pre-induction anisotropy, displayed faster mobility (Extended Data Fig. 3g–i). This observation indicates that FA-SIM not only quantitatively detects the local enhancement of crowding following phase separation but also reveals the feedforward regulatory role of the crowded environment in modulating condensate dynamics.

To further perturb intracellular crowding, we treated cytoplasmic EGFP expressing U2OS cells with trypsin, inducing cell rounding and a reduction in cytoplasmic volume. This treatment resulted in a significant increase in fluorescence anisotropy of EGFP compared to untreated controls (Fig. 3h, i), indicating elevated macromolecular crowding due to volumetric compression.

Collectively, these results establish fluorescence anisotropy of fluorescence protein as a robust reporter of macromolecular crowding and demonstrate that FA-SIM enables quantitative mapping of crowding heterogeneity and condensate dynamics at nanoscale resolution (Fig. 3j).

### FA-SIM reveals a radial crowding gradient in the microtubule network

Cells can adapt to different environments by regulating their own volume, a process that directly modulates the degree of macromolecular crowding within the cell^3, 40^. As key cytoskeleton components, microtubules (MTs) contribute to cell shape maintenance and volume remodeling^41–43^, and their dynamics are sensitive to changes in cytoplasmic viscosity and crowding, which alter the diffusion of tubulin dimers, govern polymer turnover, and influence lattice stability^17, 18, 44^. Despite the centrality of these physical parameters, conventional approaches typically manipulate crowding globally via osmotic stress that lack subcellular specificity and obscure local heterogeneities. To address this, we used FA-SIM imaging to visualized crowding-associated variations within the MT network.

FA-SIM provided markedly enhanced spatial resolution compared to FA-WF, enabling clear separation of adjacent MT filaments with distinct anisotropies that were indistinguishable under WF conditions (Fig. 4a, b). Anisotropy mapping revealed pronounced spatial heterogeneity, with higher values near the microtubule organizing center (MTOC) and lower values toward the cell periphery. To quantify this trend, we defined a centrosome-centered polar coordinate system (radial position normalized from 0 to 1) and computed mean anisotropy values within concentric bins (Fig. 4c, Extended Data Fig. 4a). This analysis revealed a graded decrease in anisotropy from the MTOC to the periphery, with mean values declining significantly from 0.22 centrally to 0.183 peripherally (Fig. 4d). Such a graded pattern is consistent with prior biophysical measurements showing higher macromolecular density and reduced molecular mobility in the pericentrosomal cytoplasm^17^. The elevated anisotropy near the cell center may result from denser MT packing or increased interactions with microtubule-associated proteins (MAPs)^22, 45^. This gradient was robust across cell types and MT probes, and persisted after fixation, where differential MT motility was eliminated while the FA gradient remained, thereby excluding MT motility as the main cause of the FA differences (Extended Data Fig. 4c, d).

**Fig. 4.**
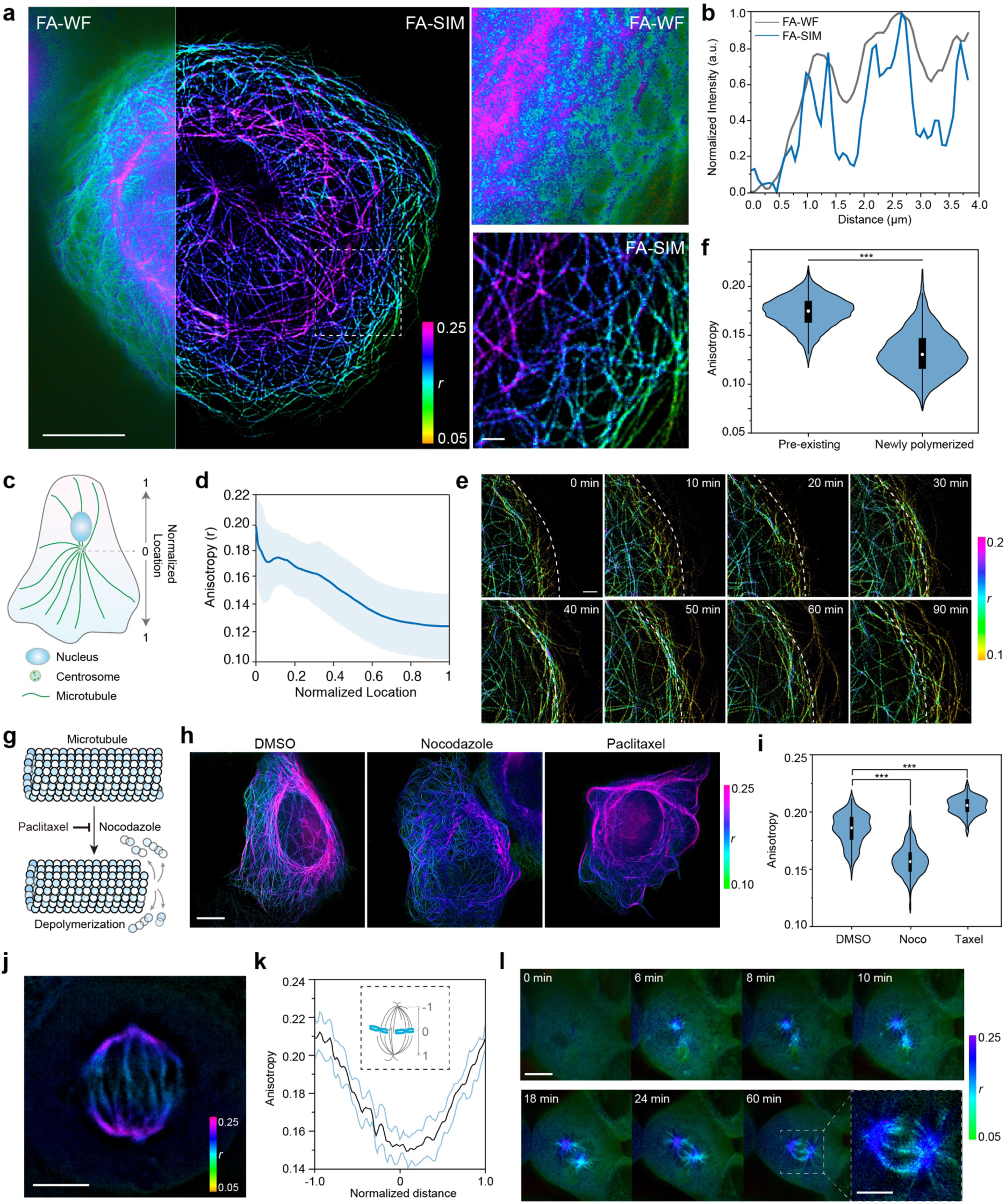
FA-SIM imaging of macromolecular crowding and MT dynamics. **a,** Representative FA-WF and FA-SIM image of MT in a U2OS cell. Scale bar, 10 μm (left) and 1 μm (right). **b,** Fluorescence intensity profiles of MTs indicated by white arrows in **(a)**. Gray: WF-FA; Blue: FA-SIM. **c,** Schematic illustration of fluorescence anisotropy quantification. The centrosome position is set as the origin (0), radiating outward to the cell edge (normalized as 1). **d,** Statistical analysis of fluorescence anisotropy across different cellular positions (0 to 1) (*n* = 20 cells). **e,** Long-term FA-SIM imaging of a U2OS cell to simultaneously monitor MT remodeling and fluorescence anisotropy dynamics. Images were acquired at 30 s intervals. The white outline indicates the initial cell boundary position at imaging onset. **f,** Quantitative comparison of FA of fluorescence proteins labeling newly polymerized or original MT in **(e)**. **g,** MT depolymerization or polymerization induced by nocodazole or paclitaxel treatment, respectively. **h,** FA-SIM images of U2OS cells treated with 0.1% DMSO (control), nocodazole, or paclitaxel. **i,** Quantitative comparison of FA of fluorescence protein for MT labeling in DMSO, nocodazole or paclitaxel treated cells. *n* = 20 cells in each group for statistic comparison. **j, k,** FA-SIM imaging of spindle in a Hela cell **(j)** and quantification of fluorescence anisotropy values between spindle poles **(k)**. (*n* = 20 cells). **l,** Long-term FA-SIM imaging of spindle remodeling in HeLa cells. Images were acquired every 30 seconds for 60 minutes. The far-right panel shows a magnified view of the fully assembled spindle. Scale bar: 1.5 μm. Error bars and error bands indicate mean ± S.D. One-way ANOVA test for the statistic calculation. *p* < 0.001: ***. Scale bars, 1 μm in **(e)**, 10 μm in **(h)**, 5 μm in **(j)** and 3 μm in **(l)**.

Time-lapse FA-SIM imaging during U2OS cell spreading further linked macromolecular crowding to MT remodeling (Fig. 4e, Supplementary Video 2). Newly polymerized MTs at the expanding periphery exhibited lower anisotropy than pre-existing MTs (Fig. 4f), consistent with reduced crowing facilitating rapid polyerization^18^. Pharmacological perturbations supported this interpretation: nocodazole treatment disrupted MTs and abolished the anisotropy gradient, whereas paclitaxel induced MT bundling and markedly increased anisotropy values (Fig. 4g–i).

Together, these results demonstrate that MT architecture and polymerization dynamics are tightly coupled to local macromolecular crowding, and FA-SIM enables quantitative, subcellular mapping of these physicochemical variations.

### Crowding-dependent organization and maturation of the mitotic spindle

Having established a radial gradient of macromolecular crowding along interphase MTs, we next asked whether analogous heterogeneity exists during mitosis, when MTs are reorganized into a highly ordered bipolar spindles. Proper spindle assembly is sensitive to cytoplasmic material properties, including viscosity, elasticity and crowding, which have been shown to influence MT nucleation efficiency, spindle size scaling and chromosome congression dynamics^46–50^. The pericentriolar material (PCM), a dense and protein-rich matrix surrounding the centrosomes, functions as the principle MTOC and has been proposed to act as a highly crowded reaction hub that concentrates tubulin dimers and regulatory factors^20, 51–53^.

Consistent with these models, FA-SIM imaging of metaphase U2OS cells revealed pronounced heterogeneity in fluorescence anisotropy within the spindle (Fig. 4j). MTs proximal to the spindle poles displayed significantly higher anisotropy than those located toward the spindle midzone, establishing a pole-oriented gradient of macromolecular crowding (Fig. 4k). This anisotropy enrichment at the poles is in agreement with recent observations that PCM undergoes mitosis-specific densification and phase-like expansion, generating a microenvironment with elevated viscosity and restricted molecular mobility^52, 54,55^. These data indicate that the centrosome-associated PCM constitutes a locally crowded microenvironment that restricts rotational diffusion of fluorophores bound to MTs.

To determine how this gradient emerges during spindle formation, we performed long-term FA-SIM imaging throughout mitotic progression (Fig. 4l, Supplementary Video 3). The time-lapse sequence captured the sequential steps of spindle assembly: tubule initially appeared as diffused cytoplasmic puncta, then accumulated at two nascent foci to form centrosomes, from which astral MTs extended radially and displayed elevated anisotropy. As astral MTs from opposing poles encountered and interdigitated, anisotropy of the MT-bound fluorescence protein increased progressively, indicating a gradual rise in local crowding accompanying spindle maturation. This dynamic evolution mirrors recent biophysical measurements showing that spindle assembly involves an increase in MT density and crosslinker occupancy, which collectively enhance the mechanical stiffness and macromolecular packing of the spindle^56–58^. FA-SIM thus not only visualizes spindle formation in real time but also uncovers the underlying microenvironmental transitions that accompany the establishment of a functional and mechanically competent metaphase spindle.

### Long-term dual-color FA-SIM enables real-time visualization of cytoskeletal interplay and crowding dynamics

Actin and MTs undergo highly coordinated remodeling during cell migration, division, and protrusion dynamics, and their spatial organization is closely associated with the local physical microenvironment of the cytoplasm^59, 60^. However, how cytoskeletal interactions correlate with, respond to, or reflect dynamic changes in the intracellular microenvironment has remained difficult to resolve with conventional fluorescence imaging. Thus, we applied dual-color FA-SIM imaging to monitor the long-term dynamics of actin-MT interactions and their relationship to the local cellular microenvironment.

Dual-color FA-SIM enabled continuous imaging for more than one hour with minimal photobleaching and negligible phototoxicity, while fluorescence intensity remained above 70% of the initial level throughout acquisition (Fig. 5a, b, Supplementary Video 4). These results demonstrate the robustness of the FA-SIM system for quantitative, time-resolved studies of organelle interactions and microenvironmental organization.

**Fig. 5.**
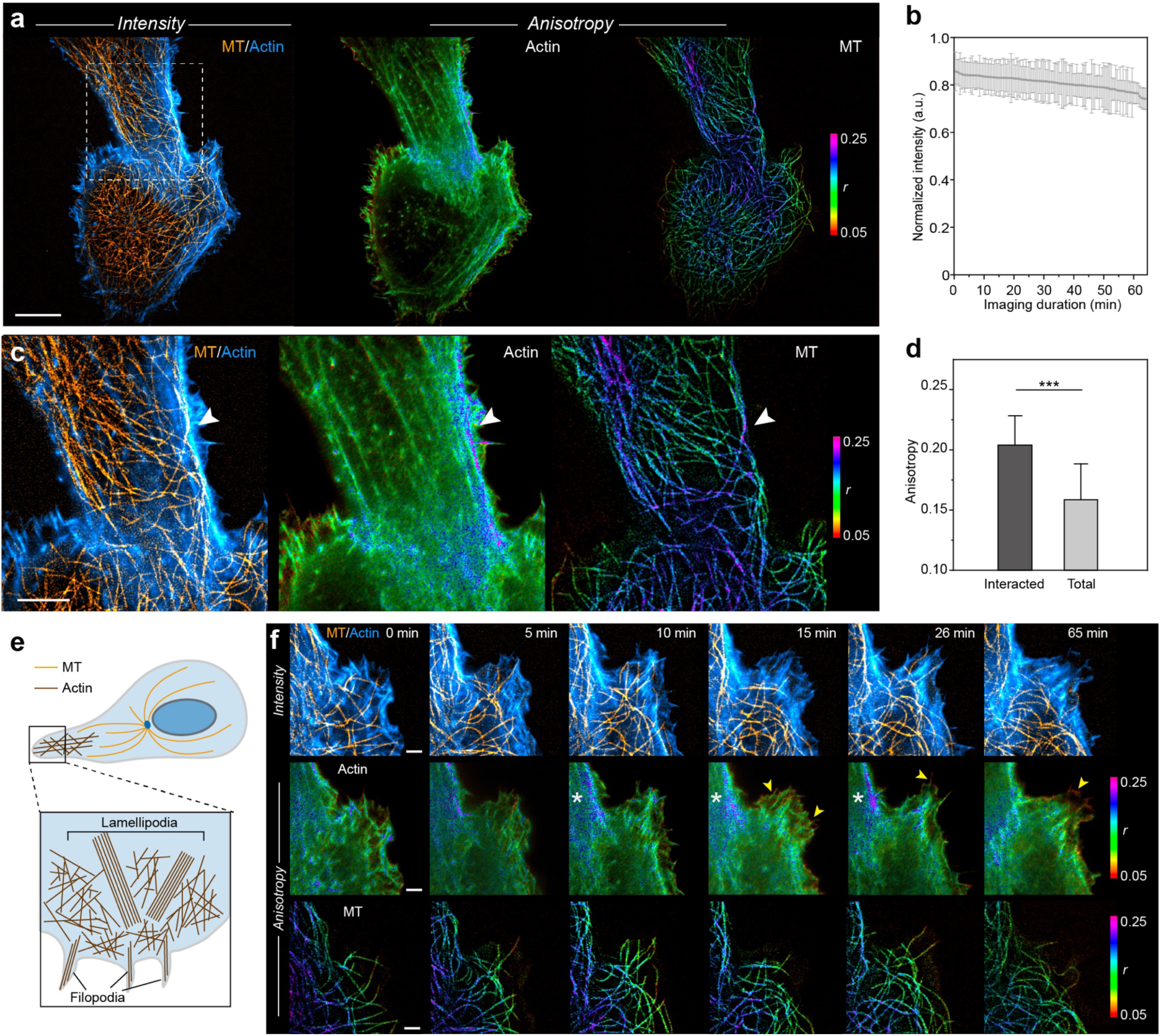
Dual-channel time-lapse FA-SIM imaging of interactions between actin and MT. **a,** Representative image of FA-SIM imaging results of actin and MT in U2OS cells. Left: merged fluorescence intensity image of actin (cyan hot) and MT (orange hot). Middle and right: FA-SIM images of actin (middle) and MT (right), respectively. **b,** Normalized fluorescence intensity during time-lapse FA-SIM imaging in **(a)**. **c,** Enlarged region within the white box in (A). The white arrows indicated the interactions between actin and MT. **d,** Quantitative comparison of FA of actin interacted with MT and total cellular actin filaments. *n* = 20 cells used for statistic comparison. **e,** Schematic illustration for lamellipodia and filopodia at cell edge. **f,** Dual-channel time-lapse interaction of actin and MT at a cell edge region. Yellow arrows indicate filopodia with lower anisotropy, and the white aster indicates lamellipodia with higher anisotropy. Images were acquired at 30 s interval for 65 min. Error bars indicate mean ± S.D. Two-tailed T-test for the statistic calculation. *p* < 0.001: ***. Scale bars, 10 μm in **(a)**, 5 μm in **(c)** and 1 μm in **(f)**.

Under these imaging conditions, we observed that actin filaments were more likely to form bundles at sites of contact with MTs, and these bundled regions displayed markedly elevated fluorescence anisotropy (Fig. 5c, d). This observation aligns with previous findings that MTs can guide or stabilize actin bundles through cross-regulatory proteins such as CLIP-170, APC, or formins, which locally modulate filament density and organization^61, 62^. The elevated anisotropy within bundles regions indicates reduced rotational freedom and increased local macromolecular crowding, providing a quantitative physical signature that complements prior qualitative descriptions of actin-MT coordination.

Long-term FA-SIM tracking further revealed distinct anisotropy signatures during protrusion dynamics (Fig. 5e). In nascent filopodia, actin underwent rapid assembly associated with consistently lower anisotropy, reflecting a more fluid and less crowded microenvironment that facilitates high-turnover assembly (Fig. 5f). This agrees with previous biophysical measurements showing that newly assembled actin filaments exhibit higher mobility and reduced packing density^63–65^. By contrast, within lamellipodia, actin frequently condensed into higher-order bundles that exhibited substantially increased anisotropy, reflecting restricted rotational dynamics associated with tighter packing and cross-linking^66–68^. This transition was reversible, as bundled actin subsequently dispersed during protrusion retraction. Throughout these cycles, MTs repeatedly invaded and remodeled protrusive domains, indicating coordinated actin-MT regulation of cell morphology. Such coordinated remodeling is consistent with established models in which MTs steer protrusion formation, deliver actin-regulating factors to the leading edge, and couple cytoskeletal reorganization to membrane dynamics^69–71^. The strong correlation between MT invasion, actin bundling, and anisotropy elevation supports the reliability of FA-SIM measurements and demonstrates that FA-SIM can capture subtle but functionally relevant microenvironment changes associated with cytoskeletal cross-talk.

## Discussion

FA-SIM addresses long-standing limitations in fluorescence anisotropy imaging by combining structured illumination with polarization-resolved detection in a manner that simultaneously enhances spatial resolution and quantitative accuracy. This integrated optical and computational framework enables quantitative FA mapping at 100-nm resolution with high temporal stability, extending FA imaging into the regime of super-resolution live-cell microscopy.

A key strength of FA-SIM is its ability to preserve quantitative fidelity while achieving long-term super-resolution imaging. This capability permits direct visualization of cellular microenvironment properties within highly dynamics subcellular structures. Through a sequence of validation experiments, including viscosity standards, nanoparticle rotation, molecular binding and perturbation assays, we demonstrate that FA-SIM provides a rigorous quantitative readout of physicochemical states.

Biological applications further illustrate the strength of this method. FA-SIM revealed nanoscale heterogeneity in cytoplasmic crowding, a radial gradient across MT arrays and pole-directed increases in spindle crowding during mitosis. These measurements capture physical features of cytoskeletal organization that were previously inaccessible with diffraction-limit FA imaging. The ability to perform hour-long, dual-color FA-SIM enabled the discovery of reversible coupling between actin-MT remodeling and local crowding changes during protrusion dynamics. These results underscore the potential of FA-based super-resolution imaging for dissecting the physical dimension of cytoskeletal regulation. Moreover, because fluorescence anisotropy sensitively reports crowding and viscosity changes induced by pharmacological perturbations, FA-SIM holds promise as a super-resolution, high-content imaging platform for profiling drug induced microenvironmental responses.

Looking forward, FA-SIM can be further combined with other microenvironment-sensitive fluorescence imaging modalities, such as fluorescence lifetime imaging^72, 73^ and microenvironment sensitive-dyes^74^ to provide multi-dimensional readouts of microenvironmental physicochemical properties. The extension of FA-SIM into three-dimensions will enable super-resolution FA mapping in 3D^75^, capturing volumetric dynamics of intracellular microenvironments.

In summary, FA-SIM microscopy bridges the domains of super-resolution imaging and quantitative biophysical measurement, offering a powerful platform to map spatial and temporal patterns of subcellular structure as well as physicochemical properties. Our findings demonstrate that cytoplasmic crowding is not a passive, uniform background, but a dynamic, structured feature of the cellular interior that is tightly coupled to cytoskeletal organization and cellular function.

## Methods

### Custom-built FA-SIM system

The FA-SIM system was constructed based on a commercial inverted fluorescence microscope (Ti2E, Nikon) equipped with a 100× objective (Nikon, CFI SR Apochromat TIRF 100× oil, NA1.49). Excitation light was provided by a laser combiner equipped with four laser lines at 405 nm (MDL-III-405-200 mW, Cnilaser), 488 nm (MDL-III-488-200 mW, Cnilaser), 561 nm (MGL-FN-561-200 mW, Cnilaser), and 639 nm (MRL-FN-639-200 mW, Cnilaser). The switching of the laser was controlled by an acousto-optic tunable filter (AOTF; AOTFnC-400.650, AA Quanta Tech), which enabled flexible selection of the excitation wavelength, as well as precise control of the laser power and exposure time according to imaging requirements. The laser output from an optical fiber (PMJ-D-460, shconnet) was collimated and reflected by a polarization beam splitter, then directed into an illumination pattern generator, which consisted of an achromatic half-wave plate, and a ferroelectric spatial light modulator (SLM; QXGA-R11, Forth Dimension Display). By adjusting the period and orientation of the grating patterns displayed on the SLM, different illumination modes could be generated. For FA-SIM imaging, two stripe orientations (0° and 90°) were employed, each comprising three phase shifts at 0, 2π/3, and 4π/3. To maximize the interference contrast of the illumination patterns, a polarization rotator composed of a liquid crystal variable retarder (LCVR; Meadowlark, LRC200) and a quarter-wave plate was used to adjust the polarization state for each illumination direction.

To ensure that the detected fluorescence polarization state remains undistorted, all mirrors in the detection light path are silver-coated planar mirrors. After passing through a polarization-maintaining dichroic mirror and tube lens, the fluorescence is extended and imaged onto a camera detector through a 4*f* imaging system (a double Föurier transform system with a total length of four focal lengths). Within the 4*f* imaging system, a polarizing beam splitter (PBS) is employed to separate *p*-polarized and *s*-polarized light, Subsequently, mirror assemblies are used to simultaneously image the two fluorescence beams with distinct polarization states onto different target areas of the camera.

### Optical characterization of FA-SIM system

The extinction ratio of the excitation light was measured at the focal plane of the objective using a rotating linear polarizer and a power meter (Supplementary Table 1).

For *G*-factor calibration, fluorescein (Sigma-Aldrich; excitation at 488 nm), Rhodamine B (Standard Imaging; excitation at 561 nm), and Cy5 (HARVEYBIO; excitation at 640 nm) were each dissolved in 0.1 M NaOH at a concentration of 10 µM.

To evaluate day-to-day reproducibility, the same dyes were separately dissolved in 0.1 M NaOH at 10 µM and then mixed in 95% glycerol (vol/vol). Solutions were sonicated to ensure complete dissolution and prepared fresh before each experiment. A 10 µL aliquot of the solution was placed between a microscope slide and a high-precision cover glass for imaging. It should be noted that system calibration was performed to ensure imaging stability and reproducibility during the imaging session, rather than to determine the absolute fluorescence anisotropy of the sample.

### Image reconstruction of FA-SIM system

The image processing was primarily performed using ImageJ software (2.0.0-rc-69/1.52p). The image reconstruction was performed using customized MATLAB software (2023a) and the code is open sourced on Github (https://github.com/PKU-Wang-Wenyi/FASIM). The detailed reconstruction steps were illustrated in **Supplementary Note 1**.

### Simulations to verify FA-SIM system

For the simulation of the FA-SIM system and verification of its defocus suppression capability in Extended Data Fig. 3c, a three-dimensional Siemens star pattern with a reference FA value of 0.3 was used as the ground truth. Based on these theoretical intensities, a three-dimensional point spread function (PSF) was simulated assuming a numerical aperture (N.A.) of 1.49 and an emission wavelength of 525 nm. Structured illumination patterns with spatial frequencies set to 0.5 times the cutoff frequency of the wide-field system were generated at orientations of 0° and 90°. The simulated intensity components were modulated by these illumination patterns and subsequently convolved with the 3D PSF to produce the synthetic raw images.

For Fig. 1f, a two-dimensional PSF was simulated with N.A. = 1.2 and an emission wavelength of 685 nm. Point-like, sinusoidal, and line-shaped structures with a spacing of 320 nm were generated as ground truth. The FA values were gradually varied from 0.1 to 0.5. Structured illumination patterns with spatial frequencies equal to 0.6 times the cutoff frequency were applied, followed by convolution with the 2D PSF. Poisson and Gaussian noise were added to obtain the final simulated raw images.

### Fluorescent beads sample preparation

Fluorescent beads with 40 nm (ThermoFisher, FluoSpheres^TM^, F8795), 100 nm (ThermoFisher, FluoSpheres^TM^, F8803) and 1000 nm (ThermoFisher, FluoSpheres^TM^, F8823) were used for FA-SIM imaging. For sample preparation, high-precision glass slides were coated with poly-L-lysine solution (poly-L-lysine/ddH₂O/PBS = 3/1/1 (vol/vol/vol)) at room temperature for at least 30 minutes to enhance the electrostatic interaction between cationic poly-L-lysine and anionic groups on the fluorescent beads. After coating, slides were rinsed twice with ultrapure water and air-dried. Carboxylate-modified microspheres were diluted in ddH_2_O at an appropriate ratio (typically 1:10,000 dilution to achieve a sparse monolayer distribution). The diluted bead solution was applied to the center of the slide. After 1 minute, the liquid was aspirated, and the slide was washed three times with ultrapure water to remove unbound beads. Finally, an anti-fade mounting medium was applied, and the prepared slides were stored overnight before imaging.

### BODIPY-biotin conjugate preparation and sample imaging

Biotin (Aladdin) was conjugated to BODIPY 493/503 NHS Ester (MREDA) as described in Supplementary Note 2. The resulting Biotin-BODIPY conjugate was dissolved in DMSO to prepare a 1 mM stock solution. 1 µM Biotin-BODIPY were mixed with varying concentrations of NeutrAvidin (Thermo Scientific) in PBS with 0.1% (vol/vol) Tween 20. All sample mixtures were kept on ice to minimize protein degradation. Prior to imaging, 10 µL of each sample was equilibrated at room temperature for 10 minutes, then placed between a microscope slide and high-precision cover glass for imaging. Each sample was imaged for 5 times.

### AZD2281-BODIPY labeling and sample imaging

AZD2281 (Macklin) was conjugated to BODIPY-FL NHS Ester (Sigma) as described in Supplementary Note 2. The resulting AZD2281-BODIPY conjugate was dissolved in DMSO to prepare a 1 mM stock solution. For live cell imaging, U2OS cells were incubated with 1 µM AZD2281-BODIPY in cell culture medium at 37°C for 1 minute. The labeling solution was then replaced with fresh culture medium prior to imaging.

### Cell culture

U2OS cells (HTB-96, ATCC) and Hela cells stably expressing tubulin-EGFP (a kind gift from Dr. Guangwei Xin at Peking University) were cultured in Dulbecco’s modified Eagle’s medium (Gibco) supplemented with 10% (v/v) heat-inactivated Fetal Bovine Serum (Thermofisher Scientific) and 1% (v/v) Penicillin-streptomycin (ThermoFisher Scientific). Cells were incubated in a sterile and humid incubator with 5% CO_2_ at 37°C.

### Live-cell imaging

Cells were plated on the 35-mm glass-bottom dishes (Cellvis, #1.5H, D35-20-1.5H) and grown typically overnight in full culture medium to reach ∼50% confluency (∼70% confluency for plasmid transfection). Before imaging acquisition, the culture medium was replaced with imaging medium containing 2% FBS in HBSS (Corning). With exceptions noted, live-cell imaging was carried out on FA-SIM system with a 100× oil-immersion objective (NA 1.49), maintained at 37°C and 5% CO_2_ via an integrated environmental chamber (Airy Technologies Co. Ltd.).

### Plasmid transfection

For plasmid transfection, cells were seeded in glass-bottom dishes (Cellvis, #1.5H, D35-20-1.5H) one day prior to imaging and cultured at 37℃ with 5% CO_2_ and 95% humidity until they reached 70% to 85% confluency. Then plasmids transfection was performed using Lipofectamine 3000 (Life Technologies). For transfection in a 35 mm glass-bottom dish: Prepare Tube A by adding 125 μL Opti-MEM, 1 μg DNA, and 2 μL P3000, then mix by pipetting up and down about 20 times. Prepare Tube B by adding 125 μL Opti-MEM and 3 μL Lipofectamine 3000, then mix by pipetting up and down about 20 times. Transfer the contents of Tube A to Tube B, mix by pipetting up and down about 20 times, and incubate at room temperature for 10 minutes. Remove the culture medium from the cells and replace it with 1.25 mL fresh Opti-MEM medium, then add the prepared transfection mixture. After 6–8 hours of transfection, remove the medium from the dish, wash twice with 1× PBS, and replace with fresh complete cell culture medium. The sample can be used for imaging after 24–48 hours.

The plasmids used in this study were as follows: EMTB-mYongHong (a kind gift from Dr. Zhifei Fu at Fujian Medical University) for MT labeling, Lifeact-mBaojin (a kind gift from Dr. Zhifei Fu at Fujian Medical University) for actin labeling, pEGFP-N1 (Addgene, 172281) for cytoplasmic EGFP expression, pmScarlet-N1 (a kind gift from Prof. Jianguo Chen at Peking University) for cytoplasmic mScarlet expression, pCDNA3.1-pCMV-PfV-GS-Sapphire (Addgene, 116933) for 40 nm genetically encoded multimeric nanoparticles expression, and ESR1 IDR-mCherry-CRY2 for light-activated phase separation.

### Light-induced phase separation and imaging

U2OS cells plated on the 35-mm glass-bottom dishes were transfected with ESR1 IDR-mCherry-CRY2 plasmid 24 to 36 h before imaging. For global activation, cells were imaged typically by use of two laser wavelengths (488 nm for CRY2 activation / 561 nm for mCherry imaging).

### FRAP experiment

FRAP experiments were performed on a Zeiss LSM980 confocal microscope equipped with a 63×/1.40 NA oil immersion objective and a temperature-controlled incubation chamber maintained at 37°C and 5% CO_2_. U2OS cells were seeded onto 35-mm glass-bottom dishes and transfected with pEGFP-N1 24 to 36 hours before imaging. Prior to imaging, the culture medium was replaced with live-cell imaging medium without phenol red.

At least three regions of interest (ROIs) with a diameter of 1.0 μm were defined with in a cell with different distance to the cell nucleus. After acquiring 5 pre-bleach frames with 488 nm laser excitation at 0.5% power, these ROIs were photobleached with 488 nm laser at 100% power for 5 iterations. After photobleaching, fluorescence recovery was immediately monitored by capturing images at 0.5 second intervals for 60 seconds using the 0.5% power.

The fluorescence intensity within the bleached ROI *I*_ROI,_ a reference background region *I*_Background,_ and a total cell region to account for total fluorescence loss *I*_Total_ were quantified using ImageJ. The data were normalized and corrected for background and bleaching during acquisition using the following equation:

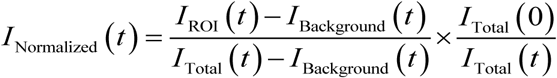

The normalized recovery curves were then plotted. 30 regions from 10 cells from three independent experiments were used for statistical comparison.

### Statistics and reproducibility

Data were expressed at mean ± S.D. in figures and text unless noted otherwise. Paired or unpaired two-tailed *t*-tests were performed as appropriate. The statistic methods and sample sizes for each experiment were listed in figure legend. All statistics were performed using Prism 10 (GraphPad). *p* > 0.05: Not significant; *p* < 0.05: *; *p* < 0.01: **; *p* < 0.001: ***.

### Data availability

An example dataset is available at https://figshare.com/s/dbc62091b1ed5d7fd6be. Any additional information about this study is available upon request from the corresponding author.

### Code availability

The original code has been deposited at Github and is publicly available at https://github.com/PKU-Wang-Wenyi/FASIM as of the date of publication.

## Acknowledgements

1. P. X. acknowledges funding support from the National Natural Science Foundation of China (62025501, 31971376, 92150301 and 62411540238) and National Key R&D Program of China (2022YFC3401100). M. Li. acknowledges the National Natural Science Foundation of China (62335008 and 62405010). We thank the National Center for Protein Sciences at Peking University in Beijing, China, for providing with PolarSIM microscope. We thank the analytical instrumentation center at Peking University for providing with FLS1000 Photoluminescence Spectrometer. We thank the Imaging Core Facility at Tsinghua University for providing with Zeiss LSM 980 imaging system. We thank the Genvivo Biotech at Nanjing, China for providing with mounting medium and other reagents.

## Author contributions

P. X. and M. Li. conceived and supervised this project. W. W. and L. Q. designed the optical path and built the FA-SIM imaging system. S. G. performed the biological experiments. W. W. and S. G. developed the FA-SIM reconstruction framework and performed data processing. S. G. composed all the figures and videos. H. W. helped in figure design. M. Liu. and Z. C. helped with BODIPY-biotin and AZD2281-BODIPY conjugation experiments. G. X. helped with biological sample preparation. Y. H. and D. K. helped with the data analysis. C. S. helped with the imaging system setup. S. G., M. Li. and P. X. wrote the manuscript with input from all authors. All authors are involved in the discussion of the results.

---

**Extended Data Fig. 1.**
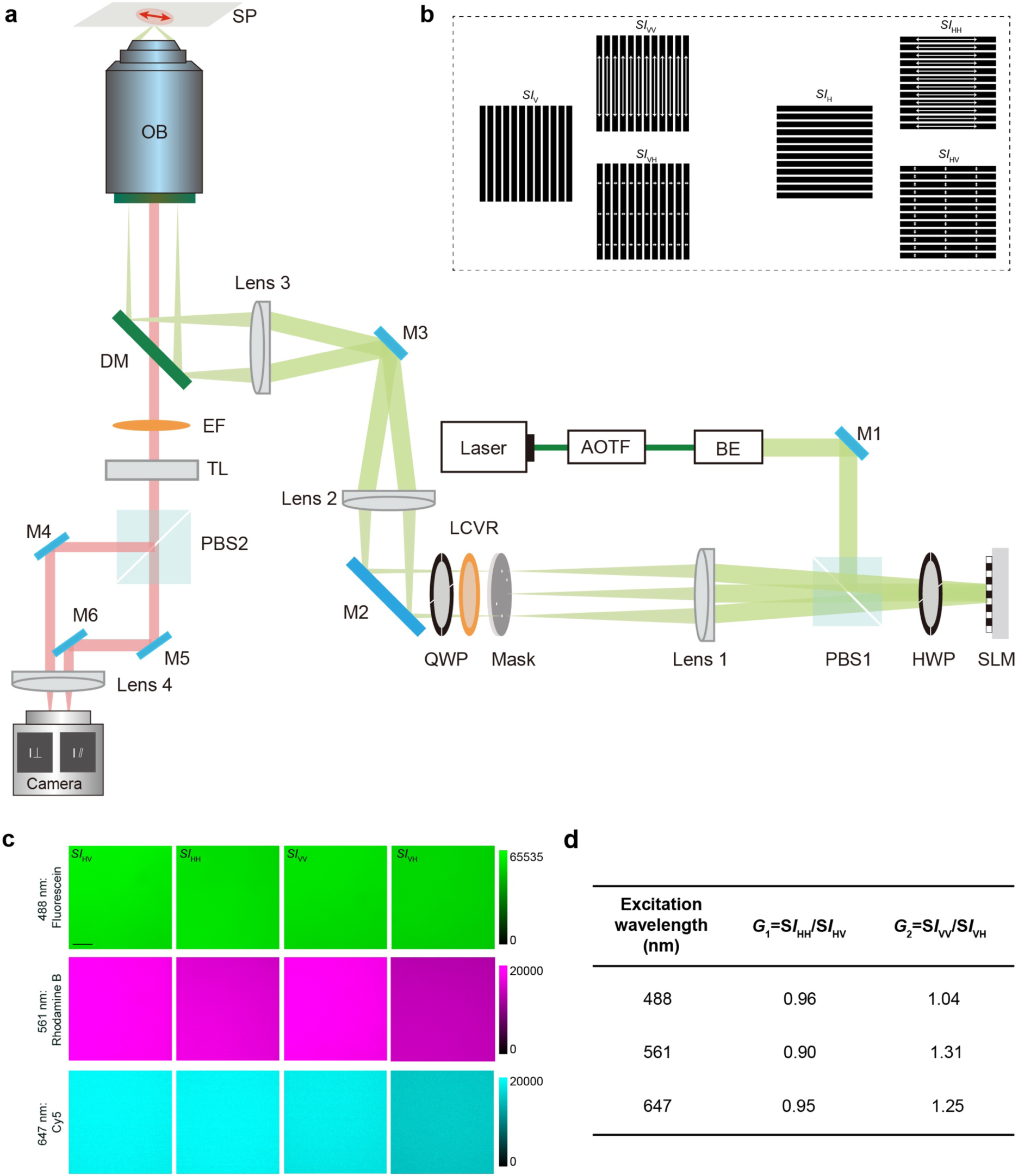
Optical setup and calibration of FA-SIM imaging system. **a,** FA-SIM imaging system optical path was composed of multiple key components: acousto-optic tunable filter (AOTF), beam expander (BE), polarization beam splitter (PBS), half-wave plate (HWP), spatial light modulator (SLM), mask, liquid crystal variable retarder (LCVR), quarter-wave plate (QWP), sample plane (SP), objective lens (OB), dichroic mirror (DM), emission filter (EF), tube lens (TL), mirrors (M1–M6), Lens1-3. **b,** Schematic diagrams of the excitation light patterns (*SI*_V_, *SI*_H_) and the corresponding camera-acquired patterns (*SI*_VV_, *SI*_VH_, *SI*_HV_, *SI*_HH_). The first subscript letter denotes the polarization direction of the excitation light, and the second indicates the polarization direction of the emitted light. The arrow directions represent the polarization orientation of the emission. **c,** Representative images of 10 μΜ solutions of fluorescein, Rhodamine B, and Cy5 in water for *G*-factor calibration. **d,** *G*-factor calibration for different excitation wavelength.

**Extended Data Fig. 2.**
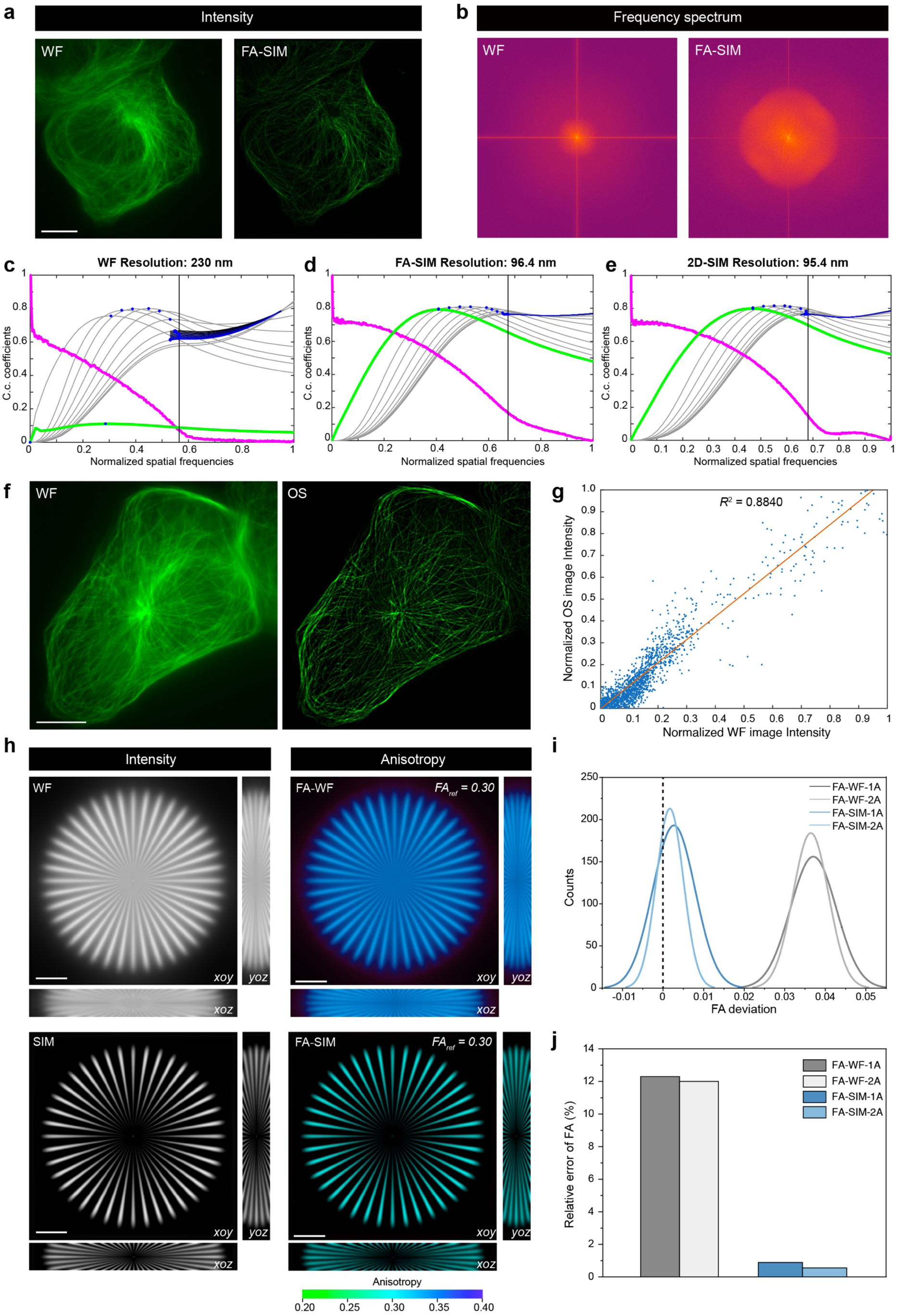
Quantitative accuracy characterization of FA-SIM imaging system. **a,** Representative fluorescence intensity images of MT in a U2OS cell under WF and FA-SIM imaging modalities. Scale bar: 10 μm. **b,** Frequency spectrum of images in (a). **c–e,** Decorrelation analysis results of WF, FA-SIM and 2D-SIM imaging systems. **f,** Wide-field (WF) and optical sectioning (OS) images of MT in a U2OS cell. Scale bars: 10 μm. **g,** Pixel intensity correlation between WF and OS images. Blue dots, raw data; orange line, linear fitted results. **h,** Simulated fluorescence intensity (left) and anisotropy (right) images of a 3D Siemens star sample (ground truth FA is 0.3), acquired with WF and FA-SIM imaging modalities. Scale bars: 10 μm. **i,** Distributions of FA deviation acquired under single angle (FA-WF-1A and FA-SIM-1A) and orthogonal dual angle (FA-WF-2A and FA-SIM-2A) imaging modalities. The dashed line indicates zero FA deviation. **j,** Relative error of FA (%) under different imaging modalities in **(h)**.

**Extended Data Fig. 3.**
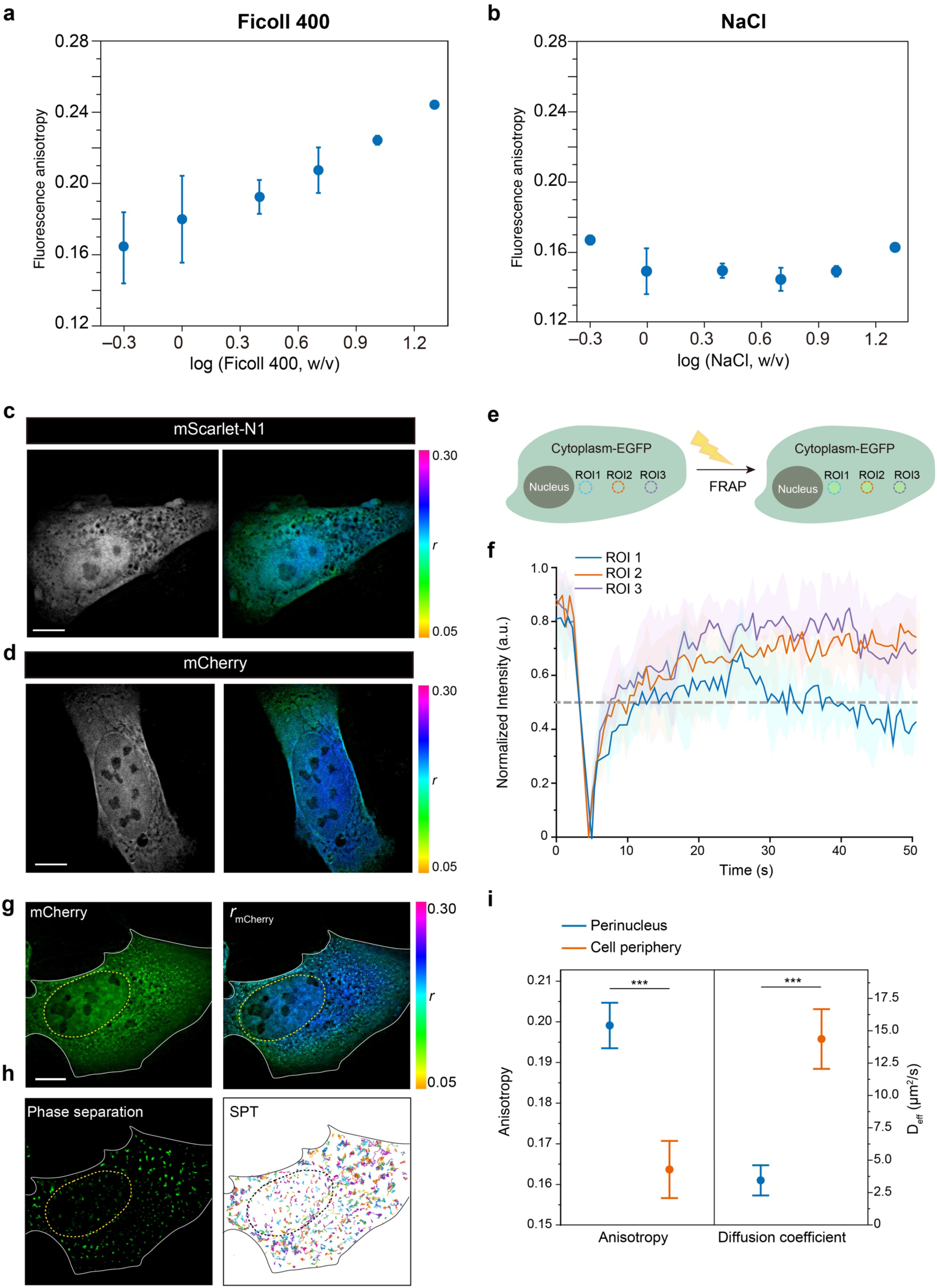
Fluorescence anisotropy of fluorescent protein serves as an indicator of macromolecular crowding. **a, b,** FA values of 1 μM EGFP in Ficoll 400 (A) and NaCl (B) solutions at varying mass fractions measured with FA-SIM imaging system. **c, d,** FA-SIM imaging of mScarlet-N1 **(c)** and mCherry **(d)** expressed in U2OS cells. **e,** Schematic of FRAP experiments in U2OS cells expressing cytoplasmic EGFP. Regions at different distances from the nucleus (ROI 1–3) were selectively photobleached with equal laser power, followed by fluorescence recovery monitoring. **f,** Fluorescence recovery curves for photobleached regions (ROI 1–3) at varying nuclear distances. Data represent 30 regions from 10 cells. Error bands indicate mean ± S.D. **g,** FA-SIM images of ESR1 IDR-mCherry-CRY2 expressing U2OS cells before phase separation. **h,** Single particle tracking (SPT) of lipid droplets after light-induced phase separation. Images were acquired at 50 Hz for 120 s. **i,** Quantitative comparison of fluorescence anisotropy (before phase separation) and diffusion coefficient of lipid droplets (after phase separation) in perinuclear and cell periphery regions. *n* = 20 cells were used for statistic comparison.

**Extended Data Fig. 4.**
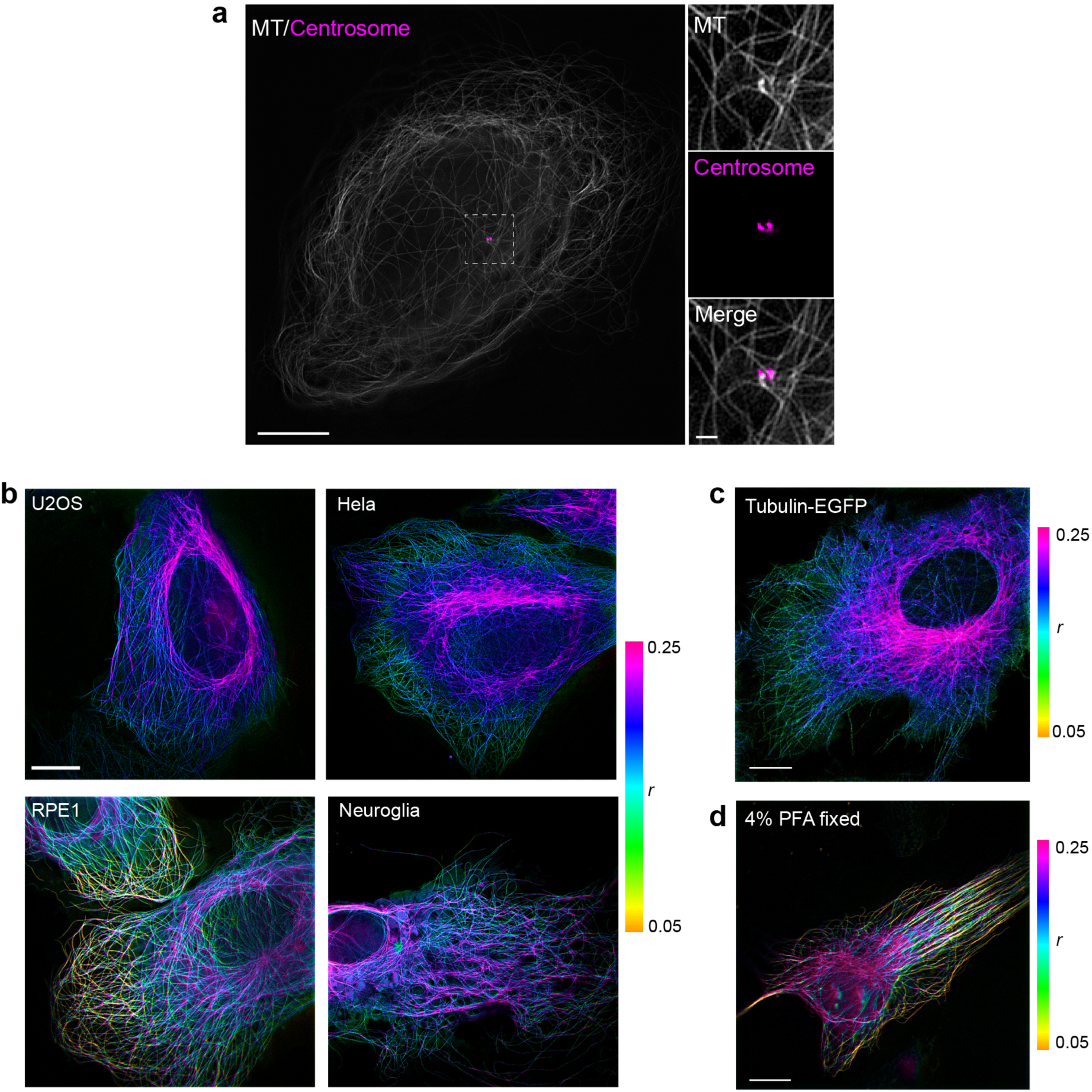
FA-SIM imaging of MT. **a,** MT labeled with EMTB-mYongHong plasmids and centrosome labeled with centrin-EGFP plasmids. Scale bar, 10 μm (left) and 1 μm (right). **b,** FA-SIM imaging of EMTB-mYongHong labeled MT in live U2OS, Hela, RPE1 and Neuroglia cells. **c,** FA-SIM imaging of Tubulin-EGFP labeled MT in live U2OS cells. **d,** FA-SIM imaging of EMTB-mYongHong labeled MT in fixed U2OS cells. Scale bars, 10 µm in (**b**, **c** and **d**).

