## Supplementary Information for "Fluorescence anisotropy structured illumination microscopy for quantitative super-resolved mapping of cell microenvironment and cytoskeletal dynamics"

[*]Contributed equally to this work

### Supplementary Notes

#### Supplementary Note 1

##### Image reconstruction of FA-SIM imaging system

**1. Sub-image extraction:** Extract the left and right sub-images corresponding to the two polarization directions from the raw image. To accomplish this, we acquire the original image under bright-field illumination and apply watershed segmentation to determine the coordinates of the two sub-images and generate corresponding masks, which were later used for the segmentation and extraction of fluorescence images.

**2. Image registration:** Register the sub-images of the left and right target planes obtained from the fluorescence channel. In this step, we imaged fluorescent beads excited at the same wavelength as the biological samples and calculate their centroid coordinates. Phase correlation–based image registration was then performed between the two sub-images to obtain an initial transformation for subsequent biological sample registration.

To align the sub-images from the transmitted ($I_{\mathrm{Trans}}$) and reflected ($I_{\mathrm{Ref}}$) channels after the polarization splitter, a two-step registration procedure was employed. First, phase correlation was used to estimate the translational offset. Denoting the Fourier transforms of the images as $F(u, v)$ and $G(u, v)$, the normalized cross-power spectrum is：

$$R(u,v) = \frac{F(u,v)G^{*}(u,v)}{\left|F(u,v)G^{*}(u,v)\right|},\ r(x,y) = \mathcal{F}^{-1}\{R(u,v)\},$$

where $G^{*}(u, v)$ is the complex conjugate of $G(u, v)$, $\mathcal{F}^{-1}$ stands for inverse Fourier transform. The peak of $r(x, y)$ gives the initial displacement $(\Delta x, \Delta y)$.

Building on the initial transformation $T$ estimated from phase correlation, intensity-based registration was subsequently applied to achieve subpixel alignment and further reduce residual registration errors:

$$\begin{bmatrix} x' \\ y' \\ 1 \end{bmatrix} = \begin{bmatrix} a_{11} & a_{12} & t_x \\ a_{21} & a_{22} & t_y \\ 0 & 0 & 1 \end{bmatrix} \begin{bmatrix} x \\ y \\ 1 \end{bmatrix}$$

$$T^{*} = \arg\max_{T} S(I_{\mathrm{Ref}}(x,y), I_{\mathrm{Trans}}(T(x,y)))$$

where $(x, y)$ are the coordinates in the transmitted sub-image $I_{\mathrm{Trans}}$, $(x',y')$ are the transformed coordinates in the reflected image $I_{\mathrm{Ref}}$, $a_{11}$, $a_{12}$, $a_{21}$, $a_{22}$ encode rotation, scaling, and shear, and

$t_x$, $t_y$ represent translations. The similarity metric $S$ quantifies the alignment quality, and the optimal transformation, $T^*$ corresponds to the parameter set that maximizes $S$.

The registration quality was quantified using the correlation coefficient (CC):

$$CC = \frac{\sum_{x,y}(I_{\text{Ref}} - \bar{I}_{\text{Ref}})(I_{\text{Trans}} - \bar{I}_{\text{Trans}})}{\sqrt{\sum_{x,y}(I_{\text{Ref}} - \bar{I}_{\text{Ref}})^2 \sum_{x,y}(I_{\text{Trans}} - \bar{I}_{\text{Trans}})^2}}$$

where $I_{\text{Ref}}(x, y)$ and $I_{\text{Trans}}(x', y')$ denote the intensity values of the reflected and transformed transmitted images at coordinates $(x, y)$ and $(x', y')$, respectively; $\bar{I}_{\text{Ref}}$ and $\bar{I}_{\text{Trans}}$ are the mean intensities of the corresponding images. A high correlation coefficient indicates successful registration and minimal misalignment between the two sub-images.

3. The masks obtained in the first step are applied to segment the biological sample images. The registration transformation derived in the second step is then applied to align the two images precisely, thereby minimizing errors in the FA calculation.

**4. *G*-factor calculation**: *G*-factor was calibrated with fluorescent dyes in aqueous solution, as shown in Extended Data Fig. 1c, d, which serves to compensate for the efficiency discrepancies between the transmission and reflection paths of the polarization beam splitter (PBS). Specifically, $G1$ is defined as $SI_{\text{HH}}/SI_{\text{HV}}$, and $G2$ is defined as $SI_{\text{VV}}/SI_{\text{VH}}$.

**5. OS-SIM reconstruction**: Optical Sectioning Structured Illumination Microscopy (OS-SIM) reconstruction is performed independently on the $SI_{\text{HH}}$, $SI_{\text{HV}}$, $SI_{\text{VH}}$, $SI_{\text{VV}}$ sub-images. By incorporating the defocused results, a more precise estimation of the FA value is achieved. The detailed processing workflow is outlined as follows:

In each sub-image, the three phase-shifted images acquired by the camera can be represented as follows:

$$I_i = I_{\text{out}}(x, y) + I_{\text{in}}(x, y)\left[1 + m\cos(2\pi k_\theta + \varphi_i\right], i = 0,1,2$$

where $I_{\text{out}}(x, y)$ and $I_{\text{in}}(x, y)$ denote the out-of-focus and in-focus components of the image, respectively. m represents the modulation depth of the illumination pattern, $k$ is the spatial frequency vector of the illumination pattern, and $\varphi_{\text{i}}$ denotes the phase of the illumination. By subtracting image pairs, the out-of-focus background can be effectively suppressed.

The basic OS-SIM reconstruction can be obtained by the root-mean-square (RMS) method[1]:

$$I_{\text{basicOS-SIM}} = \sqrt{(I_0 - I_1)^2 + (I_1 - I_2)^2 + (I_2 - I_0)^2}$$

However, the basic OS-SIM approach has several inherent limitations. Because it involves squaring operations during processing, all noise components in the image were converted into positive values, thereby reducing the signal-to-noise (SNR) ratio of the reconstructed image. Furthermore, this method requires strictly uniform phase intervals of 0°, 120° and 240°. Even minor deviations from these phase shifts can introduce residual fringe artifacts that cannot be eliminated, further degrading the image quality.

Inspired by Hilo microscopy[2], and noting that the out-of-focus background is predominantly concentrated in the low-frequency region of the image spectrum, we apply a Gaussian filter to perform low-pass filtering on the basic OS-SIM reconstruction. The Gaussian low-pass filter (GLPF) can be expressed as follows:

$$GLPF = e^{-(k_x^2 + k_y^2)/2\sigma^2}$$

here, $k_{\mathrm{x}}$ and $k_{\mathrm{y}}$ denote the spatial frequencies in the $x$ and $y$ directions, respectively, and $\sigma$ is the standard deviation of the Gaussian function, which is set to half of the fringe frequency in this context.

To recover high-frequency details, a complementary high-pass filter (GHPF) is applied to the wide-field (WF) image to extract high-frequency information, defined as:

$$GHPF = 1 - GLPF$$

Therefore, the final OS-SIM image can be obtained as follows:

$$I_{\text{OS-SIM}} = GLPF(I_{\text{basicOS-SIM}}) \times \eta + GHPF(I_{\text{WF}})$$

The scaling factor $\eta$ ensures a smooth transition between the high- and low-frequency components of the image and is related to the modulation depth of the illumination pattern.

**6. FA calculation:** After obtaining the four OS-SIM images, denoted as $I_{\text{HV OS-SIM}}$, $I_{\text{HH OS-SIM}}$, $I_{\text{VV OS-SIM}}$, and $I_{\text{VH OS-SIM}}$, the FA can be calculated using the following formula:

$$r_{\text{HH-HV}} = \frac{I_{\text{HHOS-SIM}} - G_1 \times I_{\text{HVOS-SIM}}}{I_{\text{HHOS-SIM}} + 2G_1 \times I_{\text{HVOS-SIM}}}$$

$$r_{\text{VV-VH}} = \frac{I_{\text{VVOS-SIM}} - G_2 \times I_{\text{VHOS-SIM}}}{I_{\text{VVOS-SIM}} + 2G_2 \times I_{\text{VHOS-SIM}}}$$

$$r = \frac{r_{\text{HH-HV}} + r_{\text{VV-VH}}}{2}$$

**7. Super-resolution image reconstruction.** The sub-images are combined as follows: $SI_{\mathrm{HV}}$ with $SI_{\mathrm{VV}}$, and $SI_{\mathrm{HH}}$ with $SI_{\mathrm{VH}}$, forming two illumination directions with three phase-shifted sub-images each. Using HiFi-SIM reconstruction[3], two SIM super-resolution images are obtained, which are subsequently fused to generate the final super-resolution image.

8. By integrating the FA map obtained in the fifth step with the SIM super-resolution information from this step, the FA-SIM result can be produced.

### Supplementary Note 2

#### Chemical synthesis and characterization of BODIPY-biotin and AZD2281-BODIPY

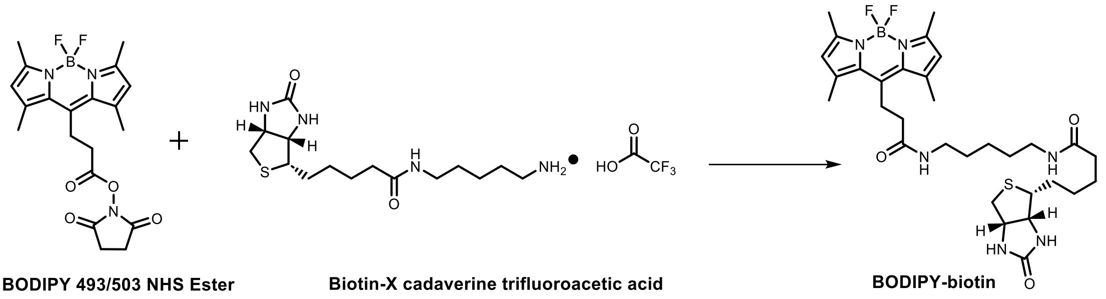

**Supplementary Fig. 1.** Chemical synthesis of BODIPY-biotin.

BODIPY-Biotin: This molecule was synthesized according to the method described previously[4]. Briefly, a mixture of BODIPY 493/503 NHS Ester (MREDA, 2.0 mg, 4.8 µmol, 1.0 eq) and Biotin-X cadaverine trifluoroacetic acid (Aladdin, 3.0 mg, 5.3 µmol, 1.1 eq) was dissolved in DMF (1.0 ml). Add *N,N*-diisopropylethylamine (DIPEA, 2.0 µL) to the solution dropwise. The resulting solution was stirred at room temperature (25 °C) for 1 h. Purification of the residue by reverse phase HPLC (eluent, a 30-min linear gradient, from 20% to 95% solvent B; flow rate, 5.0 mL/min; detection wavelength, 480 nm; eluent A (dd$H_2O$ containing 0.1% TFA (v/v)) and eluent B ($CH_3CN$)) provided BODIPY-biotin (2.1 mg, 3.3 µmol, 68% yield) as a yellow solid.

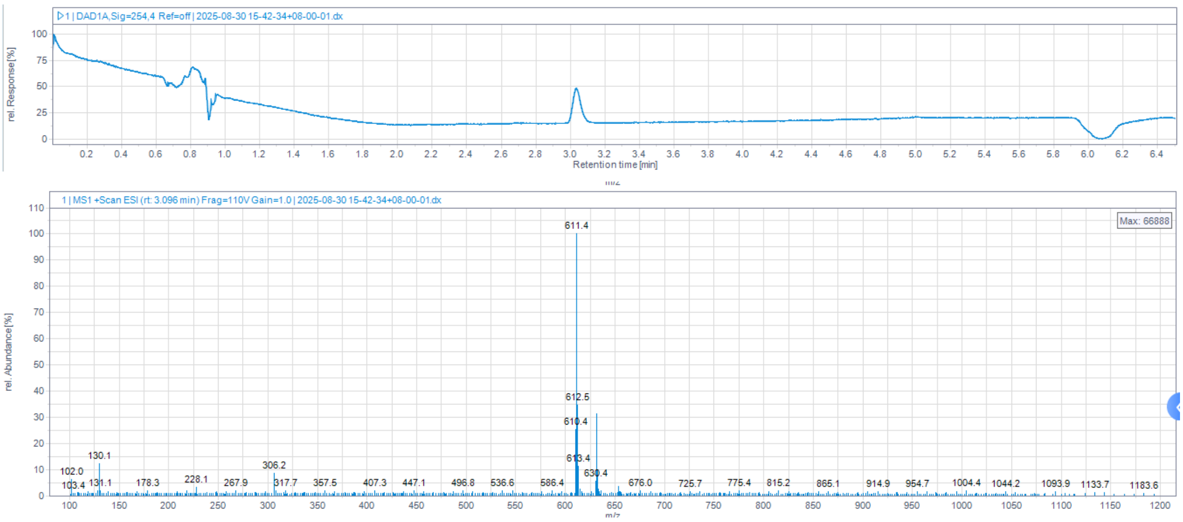

**Supplementary Fig. 2.** MS (ESI) calculated for $C_{31}H_{45}BFN_6O_3S^+$ (M-$F^+$) 611.6, observed m/z 611.4.

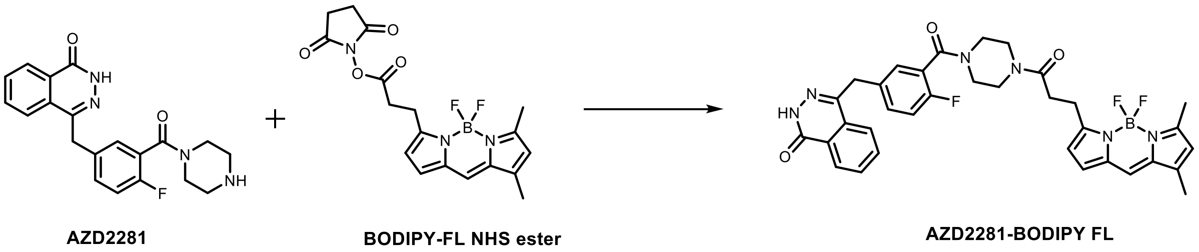

**Supplementary Fig. 3.** Chemical synthesis of AZD2281-BODIPY FL.

AZD2281-BODIPY: This molecule was synthesized according to the method described previously[4]. Briefly, a mixture of BODIPY-FL NHS ester (Sigma, 2.5 mg, 6.4 µmol, 1.0 eq) and AZD2281 (Macklin, 2.4 mg, 6.4 µmol, 1.0 eq) was dissolved in DMF (1.0 ml). Add *N,N*-diisopropylethylamine (DIPEA, 2.0 µL) to the solution dropwise. The resulting solution was stirred at room temperature (25 °C) for 1 h. Purification of the mixture by reverse phase HPLC (eluent, a 30-min linear gradient, from 20% to 95% solvent B; flow rate, 5.0 mL/min; detection wavelength, 480 nm; eluent A (dd$H_2O$ containing 0.1% TFA (v/v)) and eluent B ($CH_3CN$)) provided AZD2281-BODIPY FL (3.6 mg, 4.8 µmol, 75% yield) as a yellow solid.

**MS** (ESI) calculated for $C_{31}H_{46}BF_2N_6O_3S$ (M-$F^+$) 621.0, observed 621.4.

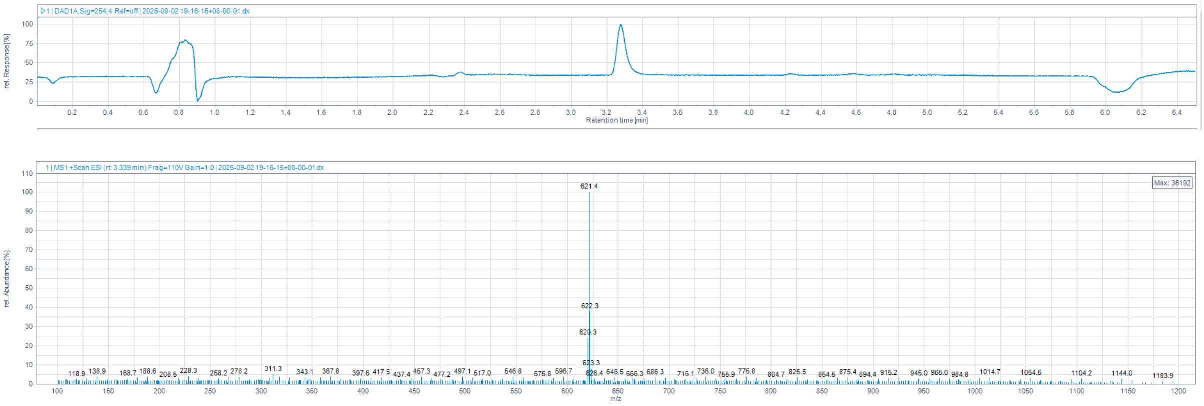

**Supplementary Fig. 4.** MS (ESI) calculated for $C_{31}H_{46}BF_2N_6O_3S$ (M-$F^+$) 621.0, observed 621.4.

### Supplementary Note 3

**Detailed methods for single-molecule tracking**

Single-molecule tracking was performed for dynamic behavior analysis in Fig. 3c and Extended Data Fig. 3h. Raw image stacks were first denoised using a combination of median filtering, Gaussian smoothing with standard deviation $\sigma = 2$, and iterative deconvolution to suppress Poisson noise and enhance contrast. Formally, each frame $I\,(x, y, t)$ was processed as:

$$I_{\text{denoise}}(x,y,t) = D(G_\sigma(\text{median}(I(x,y,t))))$$

where $G_\sigma$ represents Gaussian smoothing, median $(\cdot)$ is the median filter, and $D\,(\cdot)$ denotes RL deconvolution.

Fluorescent molecules were then detected by applying a Laplacian-of-Gaussian (LoG) filter $h_{\text{LoG}}\,(x, y)$ with scale $\sigma_{\text{LoG}}$ to highlight local intensity maxima:

$$R(x,y,t) = -(I_{\text{denoise}}(x,y,t) * h_{\text{LoG}}(x,y))$$

where * denotes 2D convolution. Candidate molecule positions were identified as local maxima exceeding an adaptive threshold $R\,(x, y, t) > \theta\,(\text{t})$, producing centroid coordinates $(x_i(t), y_i(t))$ for each frame.

Trajectories were reconstructed by linking detected positions across consecutive frames. A molecule at position $(x_i(t), y_i(t))$ was assigned to a track if its displacement from the previous frame's endpoint $(x_j(t\text{-}1),\ y_j(t\text{-}1))$ satisfied:

$$\sqrt{(x_i(t)-x_j(t-1))^2+(y_i(t)-y_j(t-1))^2} < d_{\max}$$

where $d_{\max}$ is the maximum allowed displacement.

For each trajectory, the mean squared displacement (MSD) as a function of lag time $\tau$ was computed:

$$MSD(\tau) = \left\langle \left[x(t+\tau)-x(t)\right]^2 + \left[y(t+\tau)-y(t)\right]^2 \right\rangle_t$$

where $\langle\cdot\rangle_t$ denotes averaging over all possible starting times $t$ within the trajectory. The resulting MSD curves were fitted using the generalized diffusion model:

$$MSD(\tau) = 4D\tau^\alpha$$

where $D$ is the apparent diffusion coefficient and $\alpha$ is the anomalous diffusion exponent ($\alpha = 1$ corresponds to Brownian motion). Fitting was performed in logarithmic space:

$$\log(MSD) = \log(4D) + \alpha \log(\tau)$$

followed by nonlinear regression to refine estimates of $D$ and $\alpha$.

Visualization of trajectories and MSD fitting results was performed to assess tracking quality and dynamic behavior, with each trajectory assigned a unique color for clarity.

### Supplementary Tables

#### Supplementary Table 1

**Polarization performance of FA-SIM system**

| Wavelength（nm） | Illumination direction | Maximum intensity（mW） | Minimum intensity（mW） | LCVR voltage1（V） | LCVR voltage2（V） | Depolarization ratio |
|---|---|---|---|---|---|---|
| 405 | 0° | 0.816 | 0.027 | 2.7 | 19.8 | 30.2 |
| | 90° | 0.816 | 0.019 | 4.9 | 19.6 | 42.9 |
| 488 | 0° | 9.9 | 0.301 | 1.9 | 20 | 32.9 |
| | 90° | 9.9 | 0.314 | 4 | 15 | 31.5 |
| 561 | 0° | 13.91 | 0.213 | 1.5 | 13.5 | 65.3 |
| | 90° | 13.91 | 0.339 | 3.9 | 8.8 | 41.0 |
| 640 | 0° | 5.4 | 0.106 | 2.5 | 2.8 | 50.9 |
| | 90° | 5.4 | 0.181 | 15 | 2.9 | 29.8 |

### Supplementary Videos

**Supplementary Video 1. Light-induced phase separation and FA-SIM imaging in a U2OS cell.**

Light-induced phase separation and FA-SIM imaging in a U2OS cell. Images were acquired with 100 ms exposure time and 1 s interval for 10 frames. mCherry signal was excited with 561 nm laser and phase separation was induced with 488 nm laser. Scale bar, 10 μm.

**Supplementary Video 2. Time-lapse FA-SIM imaging of microtubule remodeling in a spreading U2OS cell.**

Time-lapse FA-SIM imaging of microtubule remodeling in a spreading U2OS cell labeled with EMTB-mYongHong. Images were acquired with 100 ms exposure time and 30 s interval for 180 frames. mYongHong signal was excited with 561 nm laser. Scale bar, 10 μm.

**Supplementary Video 3. Time-lapse FA-SIM imaging of spindle remodeling in a Hela cell at metaphase.**

Time-lapse FA-SIM imaging of spindle remodeling in a Hela cell at metaphase labeled with tubulin-EGFP. Images were acquired with 100 ms exposure time and 1 min interval for 57 frames. EGFP signal was excited with 488 nm laser. Scale bar, 10 μm.

**Supplementary Video 4. Time-lapse dual-color FA-SIM imaging of cytoskeletal interactions in a U2OS cell.**

Dual-color time-lapse FA-SIM imaging of cytoskeletal interactions in a U2OS cell. Actin was labeled with Lifeact-mBaojin, and microtubule was labeled with EMTB-mYongHong. Images were acquired with 100 ms exposure time and 30 s interval for 65 min.
